# Hot and Bothered, Bees’ Gut Microbiome Shifts Under Thermal Stress and Pathogen Infection

**DOI:** 10.1101/2025.11.06.687019

**Authors:** Jennifer I. Van Wyk, Lauren Beirne, Shelly Bowder, Ellen Campbell, Milo Disharoon, Megan Dreyer, Nate Frolichstein-Appel, Aster Gill, Arreanna Jones-Ducharme, Ananda Kinkaid, Gabriel Broussard Korr, Mav McCabe, Elliot McDowell, Fernando Perez, Rio Villarreal-Mentz, Juliet Johnston

## Abstract

Understanding bumble bee gut health is imperative as these vital pollinators are subjected to pathogenic infections and thermal stress from climate change. The gut microbiome serves as an indicator for health and fitness and indicates the types of stress. To investigate the combined effects of thermal stress and pathogenic infection on bee guts, we performed a two-by-two crossed design where Bombus impatiens workers were subjected to hot conditions, pathogenic infection, or both. Incubation groups were given 2-weeks of stress conditions, with infected bees initially inoculated with Crithidia bombi, a common bee gut parasite. We measured body size, quantified the infection intensity of C. bombi using qPCR, and defined the composition of the gut microbiome using full-length 16S rRNA gene amplicon sequencing on an Oxford Nanopore Technologies Mk1D. While the core gut microbiome thrived with genera such as Bombilactobacillus and Snodgrassila which were not impacted by treatment; there were notable changes in other key organisms. Asaia bogorensis spiked in control temperature infected organisms, while species of Lactobacillus were overtaken in hot temperatures by significant increases in Apilactobacillus kunkeei. Species such as Citrobacter freundii dominated in hot infected bees suggesting an increased immunocompromised state from the combined stressors impacts on bee gut health. Our novel combined effects from thermal stress and pathogenic infection strengthen existing literature and provide new directions on how to quantify the health-state of wild bees based on their gut microbiome composition. These insights enable us to better understand how bees will be further impacted in changing landscapes.

**Importance:** We find significant changes in bees’ gut microbiome especially with an increased abundance of lactic acid bacteria. These lactic acid bacteria are often specialized: based on infection status, temperature, and the combined effects. These insights are vital to researchers, especially those studying wild bee gut health where they have an uncontrolled system and need to make assumptions about bee stress based on fitness and microbiome. Our detailed outline of relevant species will provide wild bee researchers with a baseline to determine thermal stress and recent infection status based on microbiome communities.

**Graphical Abstract:** 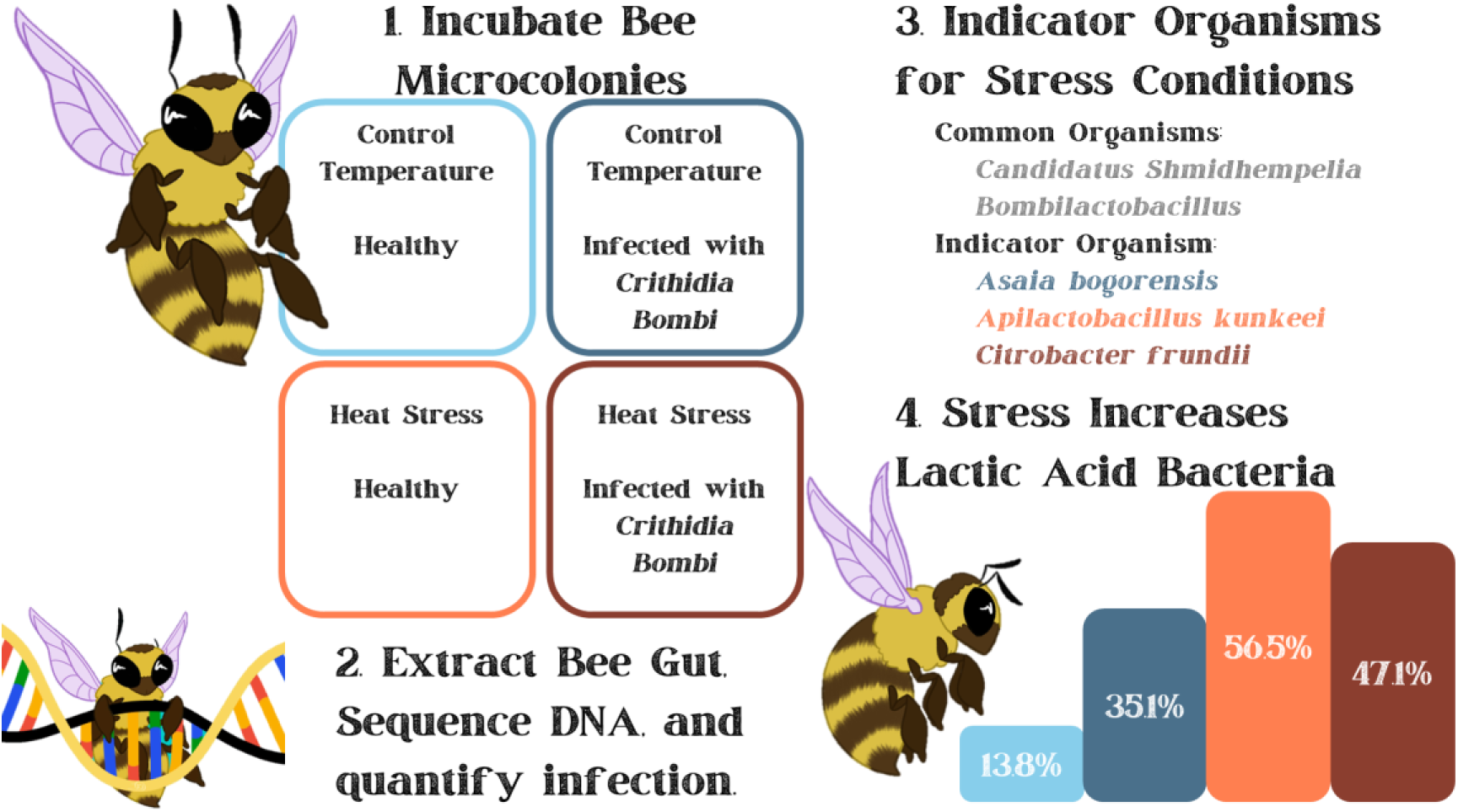

Overview of the experimental design where bees were either given control temperatures or thermal stress and/or an infection by *Crithidia* bombi. The bees microcolonies were incubated for 2-weeks before sacrificing the bees, extracting DNA, and performing downstream analysis. Full length 16S rRNA gene amplicon sequencing was performed on an Oxford Nanopore MK1D sequencer while *Crithidia* bombi infections were quantified using quantitative polymerase chain reaction.

## 2 Introduction

Bumble bees (*Bombus impatiens)* are vital pollinators in natural ecosystems and agriculture but are increasingly threatened by stressors including climate change, pathogens, pesticides, habitat loss, and declining forage quality. Comprehensive understanding of how bees respond to these pressures, and the biological mechanisms that buffer pathogen resistance, is critical for sustaining healthy populations.

Gut microbiomes are shaped through coevolution, including bees. In bees, microbes can be transmitted horizontally at flowers or vertically within colonies between nestmates^1,2^. Reviews^3–5^ have detailed their composition, diversity, and ecological drivers. Advances in 16S rRNA sequencing and the BeeBiome Consortium^4^ have enabled increasingly precise analyses linking gut health, pathogen resistance, and overall bee fitness. Recent work highlights the gut microbiome as a key mediator of these host–parasite dynamics. A core set of symbiotic genera, including *Snodgrassella*, *Gilliamella*, and *Bombilactobacillus*, support immunity and reduce infection by parasites such as *Crithidia bombi*¹, and their presence is associated with lower infection intensity and improved survival^3^.

Protective mechanisms of the microbiome include hindgut biofilms^5,6^, fermentation-derived acids that inhibit pathogens^7,8^, competitive exclusion in the nutrient-poor gut^9^, and modulation of host immune responses^10^. Studies have identified a small group of specialized bacterial taxa that may protect bees against parasites, viruses, and pathogenic bacteria^4,5,11–13^. Experimental work confirms these effects: *Bombus terrestris* workers exposed to nestmate feces developed protective gut communities and had a lower observable infection of *C. bombi*^5^, while microbial transplants in *Bombus impatiens* reduced parasite loads in individuals with higher bacterial diversity^3^. These protective symbionts are largely absent in solitary bees, suggesting that sociality facilitates beneficial microbial transmission^4,5,11,14,15^.

Environmental stressors further complicate these interactions. Bees have been shown to avoid nectar with enriched microbial communities of *Asaia astibes,* and *Apilactobacillus kunkeei*^16^ despite evidence showing *A. kunkeei* enrichment helps survivability against pathogens^17^. Both parasitism and heat stress disrupt gut microbial communities, causing dysbiosis, weakened defenses, and elevated pathogen loads^14,18^. Laboratory studies show that temperature fluctuations alter microbiome composition and infection severity, amplifying susceptibility to parasites^7,19^. While commercial bees often display greater tolerance due to consistent exposure, reduced microbial diversity in rearing environments may limit their adaptability to emerging stressors³.

Here, we test how *C. bombi* infection and heat stress interact to shape infection outcomes and gut microbiome composition in *Bombus impatiens*. Bees were incubated for two weeks before infection levels were confirmed via qPCR. Gut microbiomes were sequenced using full-length 16S rRNA amplicons, an approach not previously reported in this system, to provide species-level resolution. We hypothesized that rising temperatures alter bee gut microbiomes through thermal inactivation of *C. bombi*, offering insight into the mechanisms of pollinator resilience under climate change.

## 3 Methods

### Experimental Overview

To assess the interactive effects of pathogen infection and heat stress on the bumble bee gut microbiome, we performed a two-by-two crossed design under a controlled laboratory setting from December 9^th^ to December 23^rd^ 2024. Throughout the experiment, these four treatment groups are as follows: hot and healthy (HH), hot and infected (HI), control temperature and healthy (CH), and control temperature and infected (CI).

We randomly selected 144 female worker bumble bees (*Bombus impatiens*) from three commercially sourced colonies (Koppert Biological Systems, Howell, Michigan). Bees were individually assigned to stress treatments: heat (28 or 36 C), and pathogen (infected or uninfected control). To increase representative replication, and mimic the social environment of the eusocial insects, we housed bees communally in treatment groups of three related workers within treatment inside 500 mL plastic deli cups. Cups were placed in an incubator under complete darkness, and bees were fed 30% sucrose and pollen *ad libitum*.

### Pathogen Treatment Overview

We prepared *Crithidia bombi* inoculum from infected laboratory-maintained commercial *B. impatiens* colonies originally inoculated with parasites isolated from a wild foraging worker of the same species. We prepared a live inoculum following established methods for *Crithidia* quantification and inoculation^18^. We randomly assigned bees to pathogen or sham-inoculation treatments. Bees were individually placed in scintillation vials with perforated lids, provided 10 μL of liquid by pipette, and observed until the full volume was consumed. Only bees that consumed the full inoculum dose entered the experiment. The infection inoculum consisted of 25% sucrose solution containing 9,000 *Crithidia* cells per dose. The sham inoculum consisted of 25% sucrose in Ringer’s solution. Because infection intensity can vary among individuals even when exposed to equal inoculum doses^18^, we quantified infection at the end of the experiment using qPCR.

### Heat Treatment Overview

To create a two-by-two crossed design, the pathogen infection and control treatment groups were randomly and equally assigned between the two thermal treatments (28°C or 36°C), and maintained in incubators (VWR, 1545, 6 cubic feet, Radnor Pennsylvania, and Fisher Scientific 650D Isotemp, 5 cubic feet, Waltham, Massachusetts) for 14 days of treatment.

### Gut Microbiome Extraction

Using sterile techniques, we dissected all surviving bees to assess the gut microbiome. We removed the intestinal tract and placed them in sterile 2 mL microcentrifuge tubes, pooling replicate bees from each deli cup. For each sample, we recorded the number of bees combined and their body size. We measured the marginal wing cell length (mm) as a proxy for body size, following Nooten and Rehan^20^ (2019).

Tubes were placed in-80°C until DNA extraction was performed on March 27^th^, 2025.

### DNA Extraction

We thawed bee guts at room temperature and extracted DNA using the Qiagen DNA Blood and Tissue kit (Venlo, Netherlands) with standard protocols after lysing.

Proteinase K was added and incubated at 56°C for 3 hours prior to lyse cells before processing. Elution was performed with 100 µL of the AE Buffer to concentrate samples. DNA was promptly quantified on a NanoDrop-1000 Spectrophotometer by Thermo Scientific (Waltham, Massachusetts) to confirm purity. The average concentration of DNA extracted was 195 ± 74 ng/µL.

### Crithidia bombi quantification

*Crithidia bombi* was quantified via quantitative polymerase chain reaction from the raw DNA extracts. We used the RPB1 primers designed by Buendia-Abad et al^21^ with forward primer RPB1_F TGGTGGGTGCGATTACGAA and reverse RPB1_R TCATTGAAGATGACGTGGATAAGC. Primers were ordered from Integrated DNA Technologies (Newark, New Jersey). We used a standard qPCR protocol from New England Biolabs using their Luna Universal qPCR Master Mix (Ipswich, Massachusetts) on a QuantStudio 3 from Applied Biosystems (Waltham, Massachusetts). All samples from the infected and non-infected treatment groups were quantified. Samples with known *Crithidia bombi* infection were run in duplicate, while the uninfected samples were run singly.

### Sequencing

Full-length 16S rRNA gene amplicons were sequenced using 2 flow cells on an Oxford Nanopore Technologies MinION MK1D (Oxford, United Kingdom). Standard protocol was used based on the 16S Barcoding Kit 24 v14 with one modification where instead of utilizing the supplied magnetic beads for PCR cleanup, we used the Promega Wizard SV Gel and PCR Clean-Up System (Madison, Wisconsin). This change in PCR cleanup was due to poor yields from the supplied magnetic beads. Sequencing results yielded an average of 16,443 ± 13,303 reads per sample. Samples with low read abundances were re-run to achieve at least 5,000 reads per sample. Reads were mapped to the Silva SSU Ref NR 99 v138.2 database using nanoASV^22^. Amplicon Sequence Variants (ASVs) were then concatenated to representative species and genus levels to determine correlations across treatment groups.

### Statistics and Data Analysis

All statistical analyses were performed in RStudio v 2024.12.0 Build 467^23^. ASVs with singleton reads were removed. All samples were rarefied down to 5,000 reads per sample, and bootstrapped 1,000x to obtain a representative microbial community from each sample^24^ using the vegan 2.6.4 package in R^25–27^. Phylogenetic analysis was performed using phyloseq 1.46.0^28^. Statistical analysis for indicator species used packages bipartite 2.21^29^ and indicspecies 1.8.0^30^. Additional statistics tests utilized the dunn.test v1.3.6^31^ and ape 5.8.1^32^ packages.

## 4 Results

### Differences in Bee size and survivorship across treatments

We measured the average marginal cell length on the wings of bees, tracked survivorship, and quantified the *C. bombi* infection to determine if there were statistically significant changes based on the temperature and infection (**Figure 1**). The average marginal cell length for each group was HH = 2.55 ± 0.12 mm, HI = 2.61 ± 0.20 mm, CH = 2.65 ± 0.13 mm, and CI = 2.27 ± 0.15 mm. The control infected group (CI) was significantly smaller than all the other treatment groups (Dunn Test with Benjamini-Hochberg Correction: HH-CI p =0.012, HI-CI p = 5.01e-4, CH-CI p = 4.03e-5). The other treatment groups were not significantly different (Dunn Test with Benjamini-Hochberg Correction: HH-HI p = 0.347, HH-CH p = 0.173, HI-CH p = 0.263). There were no statistically significant differences in survivorship across any of the groups as shown in Figure 1 B&C. When quantifying the infection of *Crithidia bombi* via qPCR, three HI samples and one CI sample were below the detection limit indicating some bees were no longer infected after two weeks. Two uninfected samples initially tested positive for *Crithidia bombi* and were re-run to confirm they were false positives, confirming the infection did not impact uninfected bees. The HI group averaged 3.40 ± 2.12 log(copies/uL) while the CI group averaged 4.706 ± 1.52 log (copies/uL). There is no statistically significant difference between infections of HI and CI (Dunn Test with Benjamini-Hochberg Correction: HI-CI p = 0.1219).

**Figure 1.**
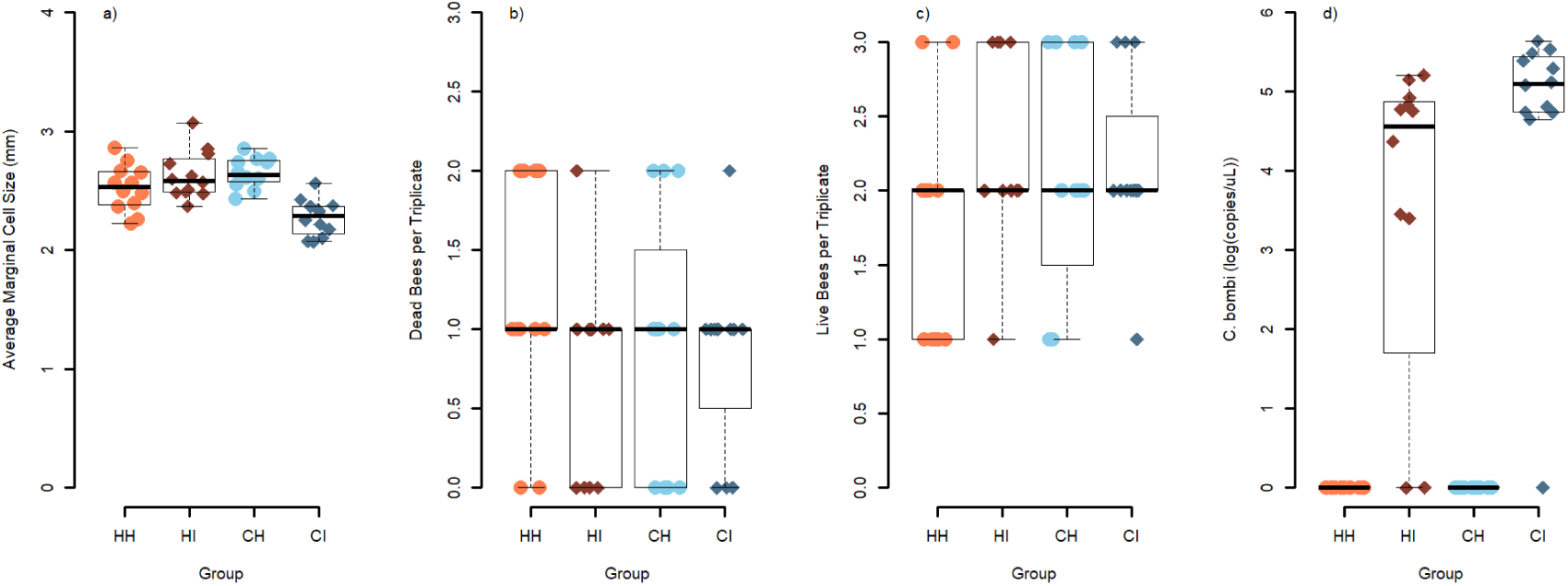
Covariates for the study were quantified for each group with a) Average Marginal Cell Length as a proxy for bee size, b) how many bees died in each incubation, c) how many bees survived in each incubation, and d) quantification of *Crithidia bombi* infections via qPCR. The HH group is shown in orange circles, the HI group is shown in dark red diamonds, CH is shown in light blue circles, and CI is shown in dark blue diamonds.

We then analyzed the trends of body size (average marginal cell length), survivorship, susceptibility to infection, and microbial diversity across the parent colonies to determine if there were major differences across the colonies (**Supplemental Figure 1**). There were no statistically significant changes based on average marginal cell length, infection quantification, and microbial diversity (Dunn Test with Benjamini-Hochberg Correction: all p > 0.05). However, parent colony B had a higher survivorship rate averaging 2.5 ± 0.61 bees/sample compared to colonies A and C which averaged 1.60 ± 0.51 bees/sample and 1.92 ± 0.64 bees/sample (Dunn Test with Benjamini-Hochberg Correction: A-B p = 2.13e-4, C-B p = 1.86e-2). There was no significant difference between colony A and colony C (Dunn Test with Benjamini-Hochberg Correction: A-C p = 0.112).

### Temperature Drives Differences in Microbial Community Diversity

Across the 48 samples, there are 1749 unique ASVs which were concatenated into 339 unique species and 39 unique genera. The average Shannon diversity index across all samples is 3.06 ± 0.96 while the average Simpson diversity index across all samples is 0.89 ± 0.07 (**Supplemental Figure 2**) at the ASV level. There were no statistically significant changes in alpha diversity based across any of the treatment groups (Kruskal-Wallis: Simpson p = 0.31 and Shannon p = 0.27).

Using the species data, we performed Bray-Curtis ordination using weighted UniFrac^33^ and NMDS to compare the microbial communities in each group (**Figure 2**). There was no statistical difference (PERMANOVA: p = 0.153) in the community based on healthy (HH and CH) and infected communities (HI and CI). The change in temperature does significantly impact on the microbial community (PERMANOVA: p = 1e-4). When analyzing both temperature and infection, there were some statistically significant differences (PERMANOVA: p = 1.6e-4). Using pairwise PERMANOVA, temperature drove differences between HH with both CH and CI (PERMANOVA with Benjamini Hochberg correction: HH-CH p = 1.8e-3, HH-CI p = 2.0e-3). Similarly, HI significantly differed from CH and CI (PERMANOVA with Benjamini Hochberg correction: HI-CH p = 6.0e-3, HH-CI p = 3.6e-2). However, there were no statistically significant differences within each temperature group based on infection (PERMANOVA with Benjamini Hochberg correction: HH-HI p = 0.29, CH-CI p = 0.27). These statistical trends followed up to the genus-level data **(Supplemental Figure 3)**.

**Figure 2.**
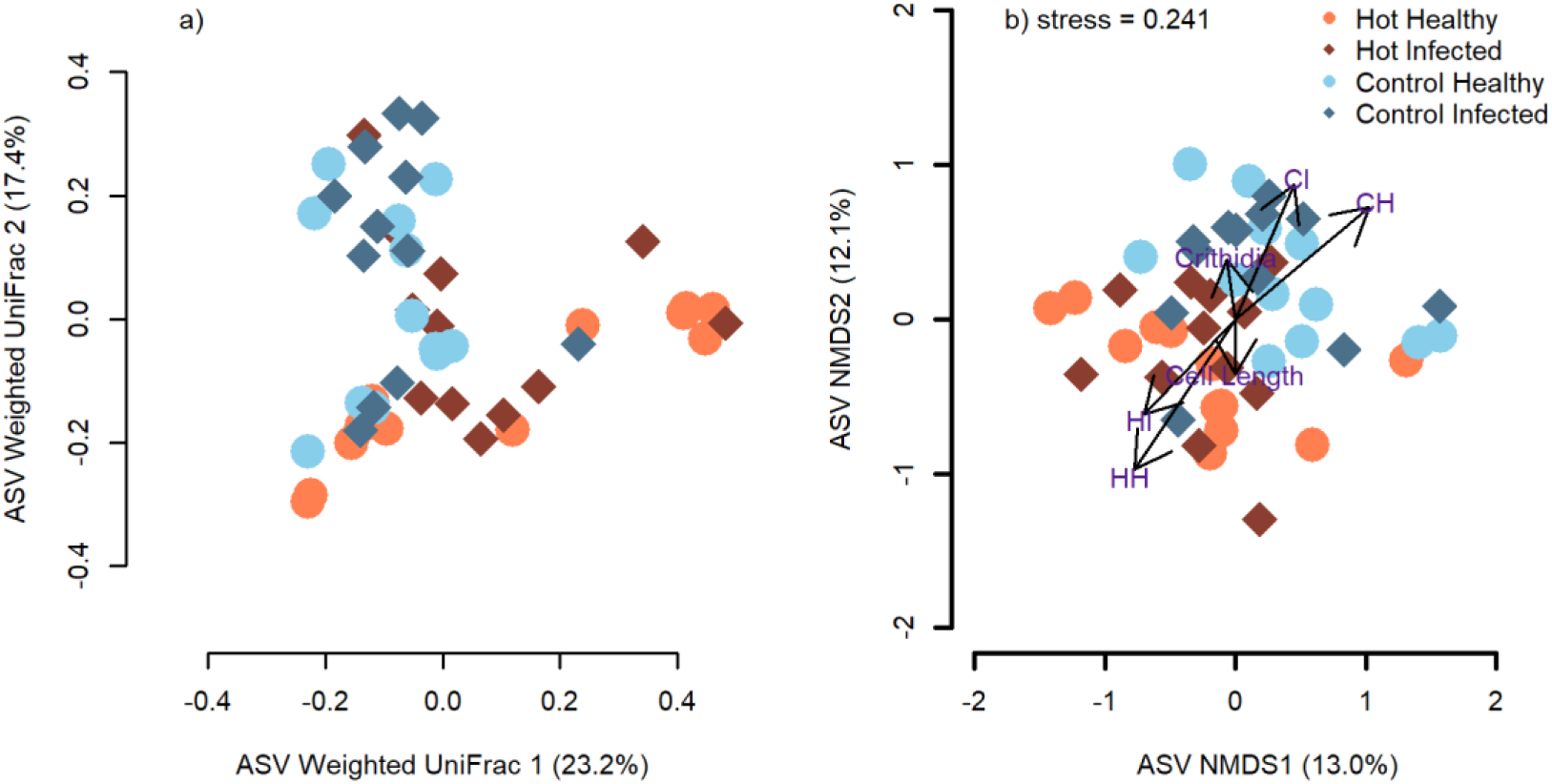
Bray-Curtis ordination was performed on a) Weighted UniFrac data from the ASV-level taxonomic assignments of the microbial community and b) NMDS. There are statistically significant differences across temperature but not based on infection. The HH group is shown in orange circles, the HI group is shown in dark red diamonds, CH is shown in light blue circles, and CI is shown in dark blue diamonds.

### Changes in Lactic Acid Bacteria

To quantify changes in the lactic acid bacteria within bees’ gut microbiomes, we explored variations across the family *Lactobacillaceae* and individual genera. There were 9 unique genera across all treatment groups within the family of *Lactobacillaceae* which included *Apilactobacillus, Lactobacillus, Bombilactobacillus, Lactiplantibacillus, Levilactobacillus, Lacticaseibacillus, Ligilactobacillus, Companilactobacilluus,* and *Lapidilactobacillus*. The family of *Lactobacillaceae* comprised an average of 56.5% ± 26.0% in the HH group, 47.1% ± 27.4% in the HI group, 13.8% ± 19.3% in the CH group, and 35.1% ± 31.2% in the CI group. All of the stressed groups were statistically significantly higher than the control CH group (Kruskal-Wallis HH-CH p = 1.7e-4, HI-CH p = 2.6e-3, CI-CH p = 0.048). A table detailing the percent abundance of each genus of lactic acid bacteria can be found in the supplemental materials (**Supplemental Table 1**).

### Indicator Species in Bee Gut Health

To determine specific organisms impacting the microbial communities, and see which are unique across groups, we summarized the entire microbial community at the genus-level **(Figure 3)** and performed multi-level pattern analysis^30^ on both the genus-level and species-level datasets. The top five genera were *Candidatus Schmidhempelia* (15.0% ± 15.9%), *Apilactobacillus* (13.8% ± 21.7%), *Lactobacillus* (12.1% ± 14.7%), *Snodgrassella* (10.5% ± 13.3%), and *Bombilactobacillus* (8.5% ± 14.4%). Out of all genera, four showed statistically significant correlations with specific groups. *Asaia* and *Neokomagataea* correlated to control temperature conditions (Multilevel Pattern Analysis: p = 5.0e-4 and p = 1.4e-2). *Acetobacter* strongly correlated with CI (Multilevel Pattern Analysis: p = 1.8e-3) and *Apolactobacillus* was prevalent in HH conditions (Multilevel Pattern Analysis: p =6.0e-4). Interestingly, we yielded different indicator organisms when treating 3 HI and 1 CI samples, which did not quantifiably yield *Crithidia bombi* infections based on qPCR, as non-infected. Truly infected CI now strongly correlated with genera *Asaia* and *Acetobacter* (Multilevel Pattern Analysis: p = 3.0e-4 and p = 1.5e-3) while truly infect HI strongly correlated to *Escherichia-Shigella, Citrobacter, Proteus,* and *Pantoea* (Multilevel Pattern Analysis: p = 3.1e-4, p = 2.4e-2, p = 3.1e-2, and p = 2.4e-2). Average percent abundance for these genus level indicator organisms are displayed in **Table 1** and **Table 2** and indicator species can be found in **Table 3** and **Table 4**. To summarize the indicator organisms, many are the species-level assignments that are members of similarly trending genera groups. This includes *Apilactobacteria kunkeei* which was more prevent in the hotter samples. Four species of *Lactobacillus,* notably *L. bombicola* which comprised a significant portion of all groups, had significantly higher abundances in CI (Multilevel Pattern Analysis: p = 0.0298).

**Figure 3.**
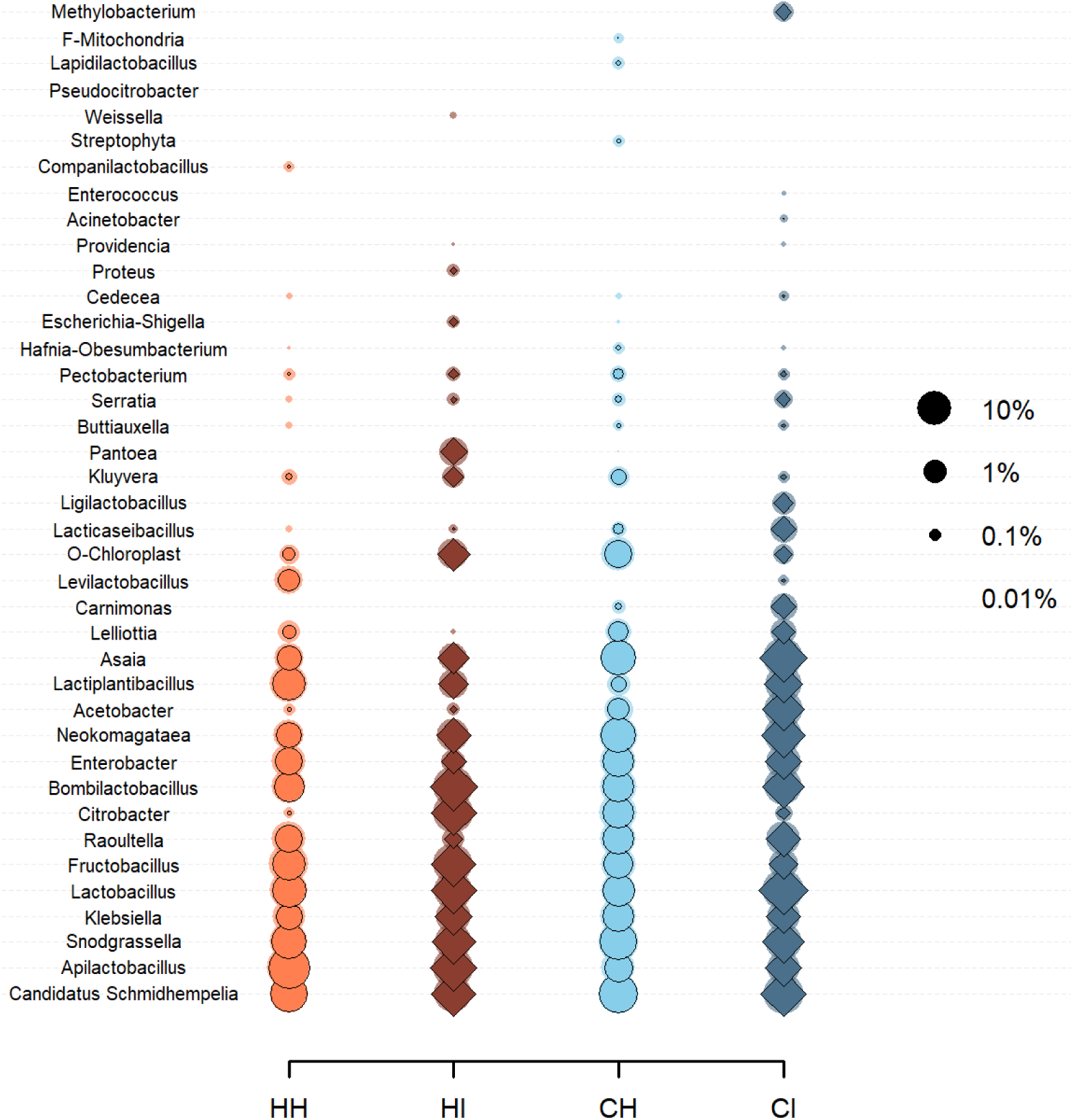
The average abundance of each community based on genus-level taxonomic identification is shown where change in size correlates to the percentage of the community, with larger sizes representing a more significant portion of the overall bacterial population. The shaded outline represents the upper standard deviation. All 12 samples per group were averaged together to identify key indicator genera across the treatment groups. The HH group is shown in orange circles, the HI group is shown in dark red diamonds, CH is shown in light blue circles, and CI is shown in dark blue diamonds.

**Table 1.** Indicator Genera based on Groups.

|  | HH | HI | CH | CI | stat | p |
| --- | --- | --- | --- | --- | --- | --- |
| *Apilactobacillus* | 34.16% | 15.17% | 2.99% | 2.91% | 0.548 | 0.0006 |
| *Acetobacter* | 0.02% | 0.03% | 0.75% | 7.55% | 0.367 | 0.0018 |
| *Asaia* | 1.47% | 1.44% | 10.29% | 17.22% | 0.589 | 0.0005 |
| *Enterobacter* | 2.20% | 0.48% | 5.15% | 2.70% | 0.426 | 0.0143 |

**Table 2.** Indicator Genera based on Quantifiable Infection.

|  | HI | CI | No Inf. | stat | p |
| --- | --- | --- | --- | --- | --- |
| *Asaia* | 1.69% | 18.48% | 5.24% | 0.615 | 0.0003 |
| *Acetobacter* | 0.04% | 8.24% | 0.33% | 0.372 | 0.0015 |
| *Escherichia-Shigella* | 0.06% | 0.00% | 0.00% | 0.467 | 0.0031 |
| *Citrobacter* | 15.73% | 0.00% | 2.13% | 0.357 | 0.024 |
| *Proteus* | 0.04% | 0.00% | 0.00% | 0.322 | 0.0305 |
| *Pantoea* | 0.89% | 0.00% | 0.00% | 0.281 | 0.0236 |

**Table 3.** Indicator Species based on Groups.

|  | HH | HI | CH | CI | stat | p |
| --- | --- | --- | --- | --- | --- | --- |
| *Lactobacillus helsingborgensis* | 0.00% | 0.07% | 0.01% | 0.24% | 0.474 | 0.0052 |
| *Lactobacillus bombicola* | 2.02% | 4.99% | 6.50% | 11.48% | 0.398 | 0.0298 |
| *Lactobacillus sp. Adhmt021* | 0.01% | 0.01% | 0.00% | 0.10% | 0.392 | 0.0311 |
| *Acetobacter orientalis* | 0.01% | 0.02% | 0.56% | 6.20% | 0.343 | 0.0058 |
| *Apilactobacillus kunkeei* | 17.65% | 10.19% | 0.75% | 1.51% | 0.417 | 0.0211 |
| *Asaia bogorensis NBRC 16594* | 0.01% | 0.02% | 0.56% | 6.20% | 0.532 | 0.0013 |
| *Asaia bogorensis* | 0.16% | 0.19% | 2.40% | 4.69% | 0.488 | 0.0006 |
| *Neokomagataea thailandica* | 1.05% | 1.07% | 7.62% | 6.53% | 0.380 | 0.0362 |
| *Asaia siamensis* | 0.00% | 0.00% | 0.07% | 0.08% | 0.378 | 0.0362 |

**Table 4.**
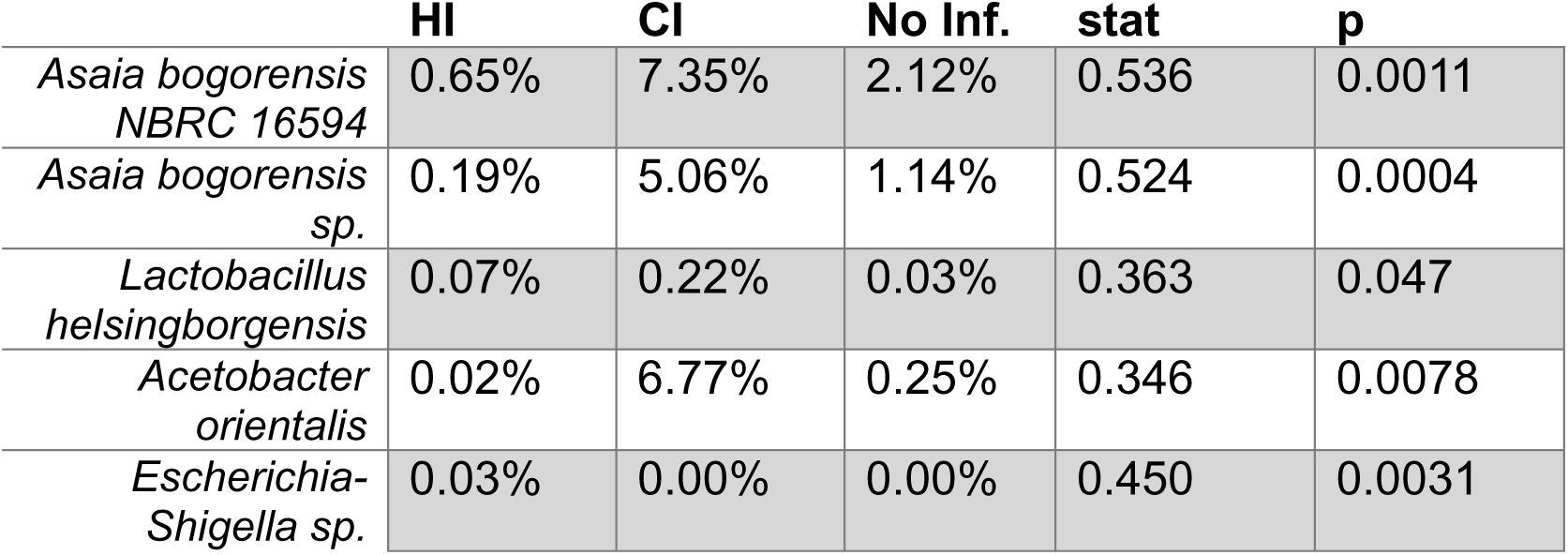
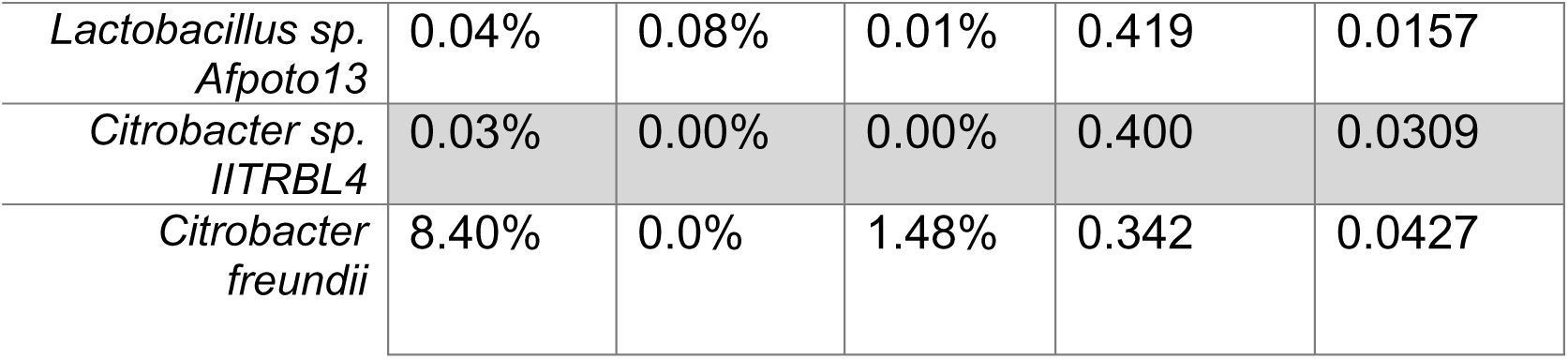
Indicator Species based on Quantifiable Infection (Top 4 for CI and HI) HI CI No Inf. stat p.

Three species of the genera *Asaia, A. borgorensis, A*. *borgorensis NBRC 16594, and A. siamensis,* were all more prevalent in the control temperature groups. These individual species with common genera drove the trends observed at the genus-level.

## 5 Discussion

### Bee gut microbiome shows resilience to infection and temperature

Overall, our bee gut microbiomes were dominated by common organisms suggesting resilience in bee gut health across thermal stress and pathogen infection. Our most abundant genera was from Candidatus *Schmidhempelia* which was previously sequenced and assembled from bumble bees and has been shown to be present in 90% of all bees^34^. Our high abundance of *Snodgrassella* is in agreement with other studies since they metabolize organic acids within the gut,^35^ and they have been shown to help influence resilience to pathogens.^36^ Another key genera was *Bombilactobacillus,* a genera that was named for its specificity to bee guts where it produces lactic acid from simple sugars and polysaccharides^37^. Similarly to *Bombilactobacillus* are the second and third most abundant genera, *Apilactobacillus* and *Lactobacillus* which significantly varied between treatment groups and are further expanded on below.

One notably absent bacterium in our dataset was *Gilliamella. Gilliamella* has been observed in bees under cold thermal stress^38^ which could explain its absence. However, it is still prevalent in many honey and bumble bee microbiomes, and is functionally important for degrading complex carbohydrates^38,39^. Additionally, most of the bees in these studies were wild bees or cultured from wild bees which are exposed to cold-shifts in the evening whereas our experimental bees were commercially reared and always maintained at constant temperatures 26 - 28°C. Since pollen sources have been shown to have little impact on bee gut diversity^40^, we believe this microbiome shift is caused by our consistently regulated temperatures. Future work to compare how commercially reared individuals respond differently to wild caught individuals is a potential area of research.

### Functional Redundancy in Lactic Acid Bacteria with Shifting Climates

Three distinct genera of lactic acid bacteria dominated the bee gut microbiome, *Apilactobacillus, Bombilactobacillus,* and *Lactobacillus.* All of these organisms were formally under the name *Lactobacillus* before rearranging them into distinct genera (i.e. past papers refer to *Apilactobacillus kunkeei* as *Lactobacillus kunkeei* and *Bombilactobacillus* was previously *Lactobacillus Firm-4*)^41,42^. Collectively these three genera averaged 48.0% of the HH community, 46.0% of the HI community, 13.4% of the CH community, and 34.4% of the CI community indicating that they are strongly associated with stress responses for bees under either thermal stress and/or infection. Species from these genera have been prevalent in other studies with thermal stress ^43^ and in bee guts infected with *Crithidia bombi*^44^ and they are commonly noted for their antimicrobial properties and disease resilience in bee guts^45^. The increased abundance of these genera outside of the control-healthy treatment group shows a strong functional redundancy where different species of lactic acid bacteria grow and respond to fulfill this necessary response when a bee is under any stress.

Our temperature treatments further show a functional redundancy in lactic acid bacteria with distinct niche preferences for several species. *Lactobacillus bombicola* thrived more in control temperatures and spiked in population during control temperature infection. This is different from past studies which noted the higher thermal tolerances of *L. bombicola* of 40°C while bee nests rarely exceed 35°C^43^. Instead, we saw an increased prevalence of *Apilactobacillus kunkeei* populations which spiked during the warmer conditions and was notably dominant In hot health (HH) groups.

While there were no statistically significant differences with *Bombilactobacillus* sp., it is notable that there were increases in infected communities (average abundance HH = 4.4%, HI = 17.5%, CH = 4.9%, and CI = 7.2%). The lack of statistically significant variations are a result of the high variation across each microcolony. We believe further replication is required or a longer incubation is necessary to promote a more stable average abundance across microcolony’s gut microbiomes.

### Bee Gut Infections in a Changing Climate

When using samples with quantifiable infection, there were several species of *Asaia* in the control infected groups, predominantly *Asaia bogorensis* which were diminished in hot infected groups. Instead, hot infected groups predominantly had various species of *Citrobacter* with *Citrobacter freundii* being the most prevalent population of a single species. *Asaia* has been reported in infected bee guts before but only as a non-core taxa^14^ and predominantly observed in field collected wild bees^46^.

Other studies reporting *Asaia* spp. observe it in mosquito guts^47,48^ and fruit flies^49^. Most *Asaia* spp. are common acetic acid fermenting organisms but our species of *Asaia bogorensis* is notably different, oxidizing acetate and lactate but not ethanol to acetic acid^50,51^. Isolates of *Asaia bogorensis* typically grow well below 30°C and are found on flowers and fruits^50,51^. Our bee’s pollen likely contains this species which could explain its significant presence in our control temperature and low abundance even in hotter temperatures, while it does not consistently show in other studies unless investigating wild bees^14,46^. It remains unclear why this species was significantly more prevalent in infected bees at control temperatures and not as prevalent in healthy control bees and could be an avenue for further research.

Bees^52,53^, beetles^54^, and even humans^55,56^ all report increased abundances of *Citrobacter freundii* when their gut microbiome is in dysbiosis. Most of the studies where bee guts are reported with *Citrobacter* observe similarly lower abundances^52,53^ (<2%) to our control temperatures and healthy bee populations. Even within our own dataset, we observed a high standard deviation where several hot infected bees had high abundances of *Citrobacter* while several reported the absence. *Citrobacter* species vary significantly with many atypical features from our most prevalent species *Citrobacter frundii*^56^. This species is most known as an infrequent cause of diarrhea in humans^55^ and can immunocompromise beetle larva^54^. This aligns with our increased abundances only in the most stressed bee population, HI, which was exposed to both thermal stress and infection.

### Survivorship Bias and Bias in Quantifying Crithidia bombi Infection

We chose to maximize the number of individual bees investigated by combining the hindguts from each microcolony, since we do not expect significant variation within the 3-bees per microcolony because they are sourced from the same parent colony.

Additionally, individual bee guts show low overall taxonomic diversity with only a handful of genera comprising a significant portion of the gut microbiome^14^. Consolidating the surviving bees from each microcolony provided us with sufficient DNA for extraction, qPCR, and also enables us to capture a better average microbiome from more replicates than individually sampling each bee.

Surprisingly there were no statistically significant changes in mortality across groups. Heat^57,58^ and *Crithidia bombi* infections^14^ have both shown to be high contributors to bee mortality. Our lower mortality is likely due to our shorter experiment which concluded within 2-weeks from infection, acclimation, and survival. This experimental design was intentional to optimize individual survival for gut microbiome analysis.

While there were no significant changes in mortality across treatment groups, three microcolonies that were infected showed up as non-detect for *C. bombi* based on qPCR. This suggests all three bees in those microcolonies survived and shed the infection^14^. The bees were only infected once during their initial feeding. Additionally, their microcolonies provided a separation screen so they were not in constant direct contact with their feces, a typical route for maintaining an infection^59,60^. To account for the lack of quantifiable infection in some bee microcolonies, we reported both indicator species from each treatment group and based on quantifiable infection as shown in Tables 1-4 since omitting the three microcolonies did impact statistical analysis.

## 6 Conclusions and Future Directions

Our research highlights that bees have a robust core microbiome that is maintained through thermal stress and pathogen stress. This core microbiome can be maintained by both individual species such as *Snodgrassella*, *Gilliamella*, and *Bombilactobacillus*, but also has functional redundancies where different species of lactic acid bacteria take over depending on temperature and stress. Finally, there are several key organisms such as *Citrobacter, Asaia,* and *Apilactobacillus* which require further research to determine if they are beneficial for bees by helping shed their pathogen through a diarrheal response for recovery, or if there is a key reason why bees with a high abundance these organisms tend to survive.

Pathogens such as *Crithidia bombi* are transmitted from bee-to-bee in fecal matter, often deposited on flora. Our experiments with microcolonies had the bees separated from their fecal matter on a mesh barrier. Future microcolony studies should explore removing any mesh barrier that separates bees from their fecal matter so bees can be in direct contact with infected feces. This will likely result in more sustained bee gut infections, where microcolonies do not have the same opportunity to recover and easily shed their infection. Additionally, because bees are social creatures and do influence each other’s microbiome^12^, upcoming research could explore the impact of bees under duress in isolation to determine if their social behaviors in larger microcolonies impacts recovery. These research directions have the potential to unravel the complex nature of bee gut microbiomes and determine a bees health status by only observing a few key indicator species in the feces. This research will help keep these vital pollinators alive and thriving despite the ongoing impacts of a warming climate and continued risk of pathogenic exposure that threaten bee colonies.

## 7 Data Availability

The sequencing data produced by this project is publicly available in the NCBI SRA under project ascension number PRJNA1305501.

## 8 Acknowledgements

This work was supported by the USDA National Institute of Food and Agriculture (award **2022-67012-38257**) to JVW; Hampshire College School for Science and Math and the Cole Science Center provided space, equipment, and facilities support. We thank Koppert Biological for supplying bumble bee colonies; the Adler lab at UMass Amherst for the Crithidia strain, lab supplies, and extensive logistical support; Sarah Steely and Willow Coville for research support; undergraduate students in course NS251 Bioinformatics and community analysis for reading previous versions of the manuscript; as well as Zoe Littlefield, June Christman, Aster Gill and Ashe Richardson-White for assistance with bee inoculations. Paid artwork of the bees was provided by Corbin Turney.

JVW and JJ conceived of the experiment, JVW executed the bee experiment, JJ executed the sequencing and conducted statistical analyses. The manuscript was written in collaboration with all authors. Undergraduate student authors are listed in alphabetical order and all contributed equally.

